# From MEG to low-density EEG: Reliable estimation of brain natural frequencies and their modulation across eyes-open and eyes-closed states

**DOI:** 10.64898/2026.04.23.720415

**Authors:** Lydia Arana, Juan José Herrera-Morueco, María Melcón, Enrique Stern, Sandra Pusil, Almudena Capilla

## Abstract

Neural oscillations are central to brain function and communication, yet they are typically characterized in terms of spectral power within predefined frequency bands, potentially obscuring their underlying functional organization. An alternative framework focuses on oscillatory frequency rather than power, revealing that each brain region exhibits a characteristic, or natural, frequency that can be estimated at the voxel level using a data-driven approach. Although this framework has been successfully applied to MEG, its broader use remains limited by cost and availability. Here, we extended this approach to EEG and validated it against MEG-derived maps, assessing its robustness across EEG channel densities (high-density, 64 channels; low-density, 32 channels) and physiological states (eyes open and closed). EEG-derived maps revealed a coherent spatial organization of natural frequencies across the cortex, reproducing the large-scale posterior-to-anterior and medial-to-lateral gradients of increasing frequency previously described with MEG. Differences between MEG and EEG were mainly confined to frontal and temporal regions, likely reflecting the differential sensitivity of the two techniques to neural source configurations, whereas posterior regions showed highly similar patterns. Importantly, this organization remained stable despite reductions in EEG sensor density and was modulated by physiological state, reproducing the well-known posterior alpha dominance during eyes-closed conditions. Together, these findings demonstrate that natural frequency mapping can be extended beyond specialized MEG research environments to low-density EEG settings, offering an accessible and scalable tool for investigating brain oscillations and their alterations in neuropsychiatric conditions.

## 1. Introduction

The human brain is constantly active, generating electromagnetic oscillations that can be captured by means of electro- and magnetoencephalography (EEG/MEG) (see Lopes da Silva, 2013, for a review). Such oscillatory activity is considered essential for brain function and information exchange (Buzsáki & Watson, 2012; Fries, 2015; Varela et al., 2001) and its alteration is closely linked to a wide range of neurological and psychiatric disorders, also referred to as “oscillopathies” (Buzsáki et al., 2013; Buzsáki & Watson, 2012; Schnitzler & Gross, 2005; Takeuchi & Berényi, 2020). Therefore, a detailed characterization of the brain’s spectral and spatial organization is key for understanding its functional architecture and identifying potential biomarkers of neuropsychiatric disorders.

Ongoing oscillatory activity has traditionally been analyzed by measuring spectral power within predefined, canonical frequency bands (i.e., delta: 0.5–4 Hz; theta: 4–8 Hz; alpha: 8–13 Hz; beta: 13–30 Hz; gamma: >30 Hz). However, frequency band boundaries are arbitrary and often result in the misclassification of similar oscillations as distinct rhythms, across species, developmental stages, individuals, and even within the same individual under different brain states (Boersma et al., 2011; Buzsáki, 2006; Buzsáki et al., 2013; Haegens et al., 2014; Mierau et al., 2017).

An alternative to the conventional approach is to focus on the precise oscillatory frequencies rather than on spectral power, thereby avoiding the use of frequency bands (Capilla et al., 2022; Donoghue et al., 2022; Frauscher et al., 2018; Groppe et al., 2013; Kalamangalam et al., 2020; Keitel & Gross, 2016; Mahjoory et al., 2020). In line with this, Capilla et al. (2022) developed a data-driven method based on clustering of power spectra, enabling identification of the natural or characteristic oscillatory frequencies of each brain region at voxel-wise resolution. Applying this approach to resting-state MEG recordings, they identified a consistent brain pattern of natural frequencies, characterized by posterior-to-anterior and medial-to-lateral gradients of increasing frequency. Specifically, natural frequencies in the alpha range predominated in posterior occipito-temporal cortices, faster beta-band oscillations were observed over frontolateral and parietal cortices, and slower delta/theta oscillations were characteristic of medial temporal and frontal areas (Capilla et al., 2022). These findings align with the natural frequencies reported in studies using single-pulse transcranial magnetic stimulation (Ferrarelli et al., 2012; Rosanova et al., 2009) and direct electrical brain stimulation (Amengual et al., 2019), as well as with frequency maps obtained via intracranial EEG (iEEG; Frauscher et al., 2018; Groppe et al., 2013; Kalamangalam et al., 2020), MEG (Keitel & Gross, 2016; Mahjoory et al., 2020), and EEG/MEG power-based approaches (Chen et al., 2008; Hillebrand et al., 2012; Janiukstyte et al., 2023; Lew et al., 2021; Mellem et al., 2017; Niso et al., 2019; Salmelin & Hari, 1994). Moreover, it has been shown that natural frequencies can be reliably obtained from individual brains, serving as a stable fingerprint that persists over the years (Arana et al., 2025). Taken together, this approach holds potential for detecting abnormalities in brain activity across the broad spectrum of oscillopathies observed in diverse neuropsychiatric conditions.

Although promising, the use of natural frequencies as a clinical biomarker based on MEG data is limited by several practical challenges, including high costs and the restricted availability of MEG scanners worldwide, which confines its application mainly to specialized research centers. In contrast, EEG offers several practical advantages, such as portability, cost-effectiveness, and broader accessibility, making it a viable option for investigating brain oscillations in both research and clinical settings (Cho et al., 2024). Its accessibility and affordability have facilitated adoption in routine clinical practice, supported by a large pool of trained professionals and ongoing technological innovations. Additionally, EEG’s flexibility to reduce the number of channels during data collection enhances efficiency, particularly in time-sensitive clinical environments. Although common EEG systems cannot fully match the spatial precision of MEG (Klamer et al., 2015; Liu et al., 2002), they provide an accessible and versatile tool that helps bridge the gap created by the global scarcity of MEG scanners.

Consequently, this study aimed to adapt the MEG-based algorithm to map the brain’s natural frequencies in EEG. To validate the approach, we compared the canonical MEG-derived map of natural frequencies with those obtained from EEG data. Furthermore, to facilitate its translation to clinical settings, we validated the method using both high- and low-density EEG (64 and 32 channels, respectively) and across the two common resting-state conditions (eyes open and eyes closed). By doing so, we sought to overcome the practical limitations of MEG and provide a scalable tool for characterizing typical brain oscillatory activity at a frequency- and voxel-wise resolution.

## 2. Materials and methods

### 2.1. Participants

Data for this study were obtained from two independent datasets: EEG recordings from the Dortmund Vital Study (Getzmann et al., 2024) and MEG recordings from the The Open MEG Archive (OMEGA; Niso et al., 2016). All procedures complied with the Declaration of Helsinki and were approved by the corresponding institutional ethics committees. Participants provided written informed consent prior to data acquisition.

#### EEG data

EEG data were obtained from the Dortmund Vital Study (Getzmann et al., 2024), comprising anonymized resting-state and task EEG recordings. Participants were selected from the Dortmund database to match the MEG group exactly in terms of age and gender. When multiple matches were available, the participant with the least artifact contamination upon visual inspection was chosen. Only two EEG-MEG pairs were excluded due to excessive artifacts, with no suitable replacement available. The final EEG group included 107 healthy volunteers (60 females, 100 right-handed, 31.1 ± 11.8 [M ± SD] years, range 20–70 years).

#### MEG data

MEG data were obtained from The Open MEG Archive (OMEGA; Niso et al., 2016), comprising anonymized resting-state MEG recordings and T1-weighted Magnetic Resonance Images (MRIs). We included MEG data from those participants in the OMEGA database who had been previously preprocessed (Capilla et al., 2022) and had a corresponding match in the EEG group based on age and gender. The final MEG group thus included 107 healthy volunteers (60 females, 100 right-handed, 31.1 ± 11.8 years, range 20–70 years).

### 2.2. Data acquisition

#### EEG data

EEG data were recorded at the Leibniz Research Centre for Working Environment and Human Factors at the Technical University Dortmund (IfADo) using a 64-channel cap based on the 10–20 system, along with a BrainVision BrainAmp DC amplifier and BrainVision Recorder software (BrainProducts GmbH). EEG signals were sampled at 1000 Hz and filtered online with a 250-Hz low-pass filter, with electrode impedances maintained below 10 kΩ. Resting-state EEG was recorded both before and after a battery of tasks, for 3 minutes with eyes closed (EC) and 3 minutes with eyes open (EO), with the EC condition acquired first. Only the resting-state EEG (EO and EC) recorded prior to the tasks were included in the present study.

#### MEG data

MEG data were recorded in a magnetically shielded room at the Montreal Neurological Institute (MNI, McGill University) using a whole-head CTF MEG system with 275 axial gradiometers and 26 reference sensors at a 2400-Hz sampling rate. An online 600-Hz low-pass anti-aliasing filter and CTF third-order gradient compensation were applied. Participants remained awake with eyes open (EO), fixating on a cross for 5 minutes during the recording.

### 2.3. Preprocessing

#### EEG data

For EEG data preprocessing, we adapted the automatic pipeline DISCOVER-EEG (Gil Ávila et al., 2023). Both EO and EC runs were preprocessed separately following these steps. First, the data were downsampled to 512 Hz to reduce computational load. The power line artifact at 50 Hz (and harmonics at 100 and 150 Hz) was reduced by means of spectrum interpolation (Leske & Dalal, 2019), followed by high-pass filtering with a default transition band of 0.25–0.75 Hz. Channels were marked as artifactual and removed from the data if they met any of the following criteria (1) being flat for more than 5 seconds, (2) having a z-scored noise-to-signal ratio greater than 4, or (3) showing a correlation with nearby channels lower than 0.8 (default parameters in Gil Ávila et al., 2023; Pernet et al., 2021). Then, the data were rereferenced to the average reference.

Subsequently, five repetitions of the following steps were performed. (1) Artifacts were removed using independent component analysis (ICA) with the ICLabel algorithm, which automatically labels components and classifies them into distinct categories (Pion-Tonachini et al., 2019); components with a probability higher than 80% of being classified as “Muscle” or “Eye” were rejected (Gil Ávila et al., 2023; Pernet et al., 2021). (2) Previously removed channels were interpolated using spherical splines (Perrin et al., 1989). (3) Bad time segments containing artifacts were marked but not removed to avoid introducing edge effects in the continuous data. Originally, the DISCOVER-EEG pipeline removed bad segments using the Artifact Subspace Reconstruction (ASR) method (Mullen et al., 2015), which identifies segments with abnormally high power. In our approach, we retained the original continuous data and stored a mask of artifactual segments to exclude them from subsequent analysis. The parameters for identifying bad segments, based on Pernet et al. (2021) and Gil Ávila et al. (2023), were adapted to suit our data. Specifically, segments that deviated from the calibration data by a variance 60 times higher were marked as bad (compared to the default value of 20, which led to the removal of alpha waves in our data), and segments containing 10% or more artifactual channels were also marked as bad (a stricter criterion than the default 25%, ensuring that remaining artifacts were detected). The maximum tolerance for standard deviations was set to 9. Among the five repetitions of these three steps, the iteration whose bad time segment mask was closest to the mean across repetitions was selected as the final cleaned dataset. As a final sanity check, the cleaned data were visually inspected to confirm successful artifact removal.

The number of bad channels interpolated was 2.3 ± 1.9 for EO and 2.4 ± 1.7 for EC. The number of eye components rejected was 1.9 ± 0.6 for EO and 1.4 ± 0.9 for EC. Finally, the number of muscle components rejected was 2.9 ± 3.3 for EO and 1.9 ± 2.5 for EC.

#### MEG data

We employed MEG data preprocessed as described in Capilla et al. (2022). Briefly, the MEG signal was denoised using principal component analysis (PCA), high-pass filtered at 0.05 Hz, and corrected for power line artifacts via spectrum interpolation (Leske & Dalal, 2019). The signal was then demeaned, detrended, resampled at 512 Hz, and corrected using ICA. Data segments containing residual artifacts were marked and excluded from further analysis. To match the duration of the EEG recordings, the 5-minute MEG sessions were truncated to 3 minutes.

### 2.4. Brain maps of natural frequencies

Data processing was conducted using FieldTrip (version 20230118; Oostenveld et al., 2011) and custom in-house Matlab code. The scripts necessary to reproduce all the analyses are available at https://github.com/necog-UAM.

The procedure began with the extraction of individual natural frequency maps for each participant. EEG/MEG signals were reconstructed in source space, and power spectra were computed for each voxel and participant, followed by clustering using k-means. The most characteristic oscillatory frequency of each voxel was then identified to generate individual natural frequency maps for MEG (where only eyes-open recordings were available, MEG-EO), as well as for high- (EEG-64-EO) and low-density EEG (EEG-32-EO), including both eyes-open (EEG-64-EO) and eyes-closed (EEG-64-EC) conditions. Subsequently, these maps were compared across recording modalities (MEG-EO vs. EEG-64-EO vs. EEG-32-EO) and eye conditions (EEG-64-EO vs. EEG-64-EC) to assess their similarity and to identify regions exhibiting significant differences. The following sections describe each step of the process in greater detail.

#### 2.4.1. Reconstruction of source-level time series

##### EEG data

Prior to source reconstruction, an offline channel selection was performed, reducing the 64 recorded channels to a 32-channel subset in the EO condition to represent high- and low-density EEG configurations, respectively. This selection was based on the channels available in the 64- and 32-channel caps compatible with the recording system used in Getzmann et al. (2024) (BrainVision BrainAmp, BrainProducts GmbH) and followed the 10-20 system for channel coordinates and labeling (Klem et al., 1999). As a final preparation step before source reconstruction, both high- and low-density EEG data were rereferenced to the average reference.

EEG source reconstruction was performed on the standard Montreal Neurological Institute (MNI) brain. To generate the head model, the MNI MRI (Holmes et al., 1998) was co-registered to the EEG standard 10–05 electrode positions (Oostenveld et al., 2003; Oostenveld & Praamstra, 2001), selecting the corresponding EEG channels in our data (i.e., 64 or 32 channels). A standard boundary element method (BEM) volume conduction model was employed (Oostenveld et al., 2003), segmented into 1 cm3 voxels. Following Capilla et al. (2022), voxels were restricted to the cortical surface, excluding non-cortical regions such as the cerebellum and subcortical structures, resulting in 1925 voxels. Four dipoles overlapped with electrode locations and were therefore discarded, yielding a final set of 1921 voxels. Lead fields were subsequently computed for each voxel. Finally, to address the central bias of beamformer source estimates, whereby reconstructed activity is stronger in deeper regions than in cortical areas (Shapira Lots et al., 2016), we normalized the source-space signal of each voxel by dividing it by its standard deviation across time.

##### MEG data

Source reconstruction for MEG was performed on each participant’s individual MRI. T1-weighted MRIs were co-registered with the MEG coordinate system using a semi-automatic procedure based on a modified version of the Iterative Closest Point algorithm (Besl & McKay, 1992). The forward model was obtained using a realistic single shell volume conductor model (Nolte, 2003). A standard MNI 3D grid (1 cm3 resolution) was adapted to each individual’s brain volume, excluding regions outside the cerebral cortex and hippocampus, leaving 1925 voxels for analysis. Lead fields were then computed for each grid point.

From this step onward, all analyses were conducted using the same procedures for both EEG and MEG. Source-level time series were reconstructed using linearly constrained minimum variance (LCMV) beamforming (Van Veen et al., 1997). The spatial filter weights were derived from the covariance of the artifact-free data, with the regularization parameter lambda set to 10% and lead fields normalized to mitigate depth bias. These beamforming weights were then applied to the sensor-level data to estimate source-space time series. To mitigate the center-of-head bias (Shapira Lots et al., 2016), the source-space signal of each voxel was normalized by dividing it by its standard deviation across time.

#### 2.4.2. Frequency analysis of source-level data

A Hanning-tapered sliding window Fourier transform was applied to the source-reconstructed signals with a step size of 200 ms. To enhance the quality of single-subject natural frequency maps (Arana et al., 2025), spectral analysis was conducted at a higher temporal resolution than in Capilla et al. (2022) (200 ms vs. 500 ms steps). Power was estimated across 61 logarithmically spaced frequency bins ranging from 1.7 to 34.5 Hz, as previous findings indicate that higher frequencies primarily reflect artifactual activity at rest (Capilla et al., 2022). The sliding window length was frequency-dependent and set to five cycles per frequency bin, attenuating the 1/f aperiodic component. To prevent edge effects, 5.8-s intervals (corresponding to 10 cycles at the lowest frequency of 1.7 Hz) at the beginning and end of each artifact-free segment were excluded from the analysis. Thus, we obtained a set of 785 ± 88 power spectra per voxel, session, and participant for EEG-64-EO and EEG-32-EO, 799 ± 111 spectra for EEG-64-EC, and 759 ± 88 spectra for MEG-EO. Finally, each power spectrum was normalized by its total power across frequencies to obtain relative power values, ensuring that all power spectra are on the same scale.

#### 2.4.3. Whole-group cluster analysis of power spectra

To identify different patterns of source-reconstructed oscillatory activity, we performed a whole-group k-means clustering using the cosine as the distance metric to emphasize differences in the shape of the power spectra rather than their amplitude (Keitel & Gross, 2016). Clustering was conducted separately for MEG-EO, EEG-64-EO, EEG-32-EO, and EEG-64-EC, randomly selecting 252 power spectra per voxel and participant from each condition (corresponding to the number of spectra from the participant with the fewest available spectra across all conditions). Thus, we introduced a total of 252 power spectra x 1925 voxels x 107 participants for MEG and 252 power spectra x 1921 voxels x 107 for EEG. The number of clusters was set to 25, and clustering was performed over 5 runs with a maximum of 200 iterations each to optimize results. The final solution was selected based on the lowest total within-cluster distance (see Capilla et al., 2022). Cluster centroids for MEG-EO, EEG-64-EO, EEG-32-EO, and EEG-64-EC are displayed in Supplementary Material (Supplementary Figures 1, 2, 3, and 4, respectively).

#### 2.4.4. Single-subject brain maps of natural frequencies

After training the k-means clustering model on the whole-group data, we generated single-subject maps of natural frequencies (Arana et al., 2025), producing one map per individual and condition (MEG-EO, EEG-64-EO, EEG-32-EO, and EEG-64-EC). For each participant and voxel, all precomputed power spectra were assigned to the whole-group clusters by computing the distance between each spectrum and the cluster centroids, with each spectrum allocated to the cluster with the nearest centroid (i.e., smallest cosine distance). To characterize the oscillatory frequency of each cluster, the frequency peak of its centroid was identified, discarding centroids without detectable peaks, as spectral peaks are required to confirm the presence of genuine oscillatory activity (Donoghue et al., 2022). When two peaks were detected, we assessed whether they formed a harmonic pair (e.g., 10 and 20 Hz), allowing for a ±1 Hz tolerance. In such cases, only the lower (fundamental) frequency was considered for subsequent analyses. Alternatively, if the peaks were not harmonically related (e.g., 7 and 22 Hz), both were considered equally, each contributing 50%.

Next, we calculated the proportion of power spectra assigned to each cluster and normalized these values across voxels using z-scores. To increase frequency resolution, centroid peak frequencies and their corresponding z-values were interpolated tenfold. The oscillatory frequency associated with the highest z-value was then provisionally assigned as the voxel’s natural frequency. Finally, to account for abrupt transitions between neighboring voxels (e.g., delta to beta), spatial smoothing was applied across neighboring voxels (Arana et al., 2025). For each voxel, neighbors within a sphere of < 1.5 cm radius were identified, and a t-test was conducted across these surrounding voxels for each frequency bin. The frequency with the highest t-value was then assigned as the voxel’s final natural frequency. Voxels for which the t-value was not statistically significant (*p* > 0.05), indicating an unstable natural frequency across neighbors, were assigned a missing value.

#### 2.4.5. Group-level brain maps of natural frequencies

After computing the single-subject maps of natural frequencies, we generated the group-level maps for each condition. For each voxel, we calculated the distribution of natural frequencies across participants. These distributions were represented as smoothed histograms using logarithmically spaced frequency bins to ensure appropriate resolution across the entire frequency range. The most prominent peaks were then identified as candidate oscillatory modes and subsequently characterized by fitting Gaussian functions. The Gaussian peak with the highest amplitude was selected, and its corresponding frequency value was assigned as the voxel’s natural frequency. This procedure effectively identified the oscillatory frequency most consistently observed across participants, corresponding to the most prevalent value for that voxel.

### 2.5. Similarities in natural frequencies across recording modalities

Following previous work that established the robustness of natural frequencies using MEG (Arana et al., 2025; Capilla et al., 2022), we aimed to compare the correlation between the group-level natural frequency maps obtained from EEG and MEG. To this end, we represented the group-level natural frequency as vectors containing one frequency value per voxel, yielding 1925 values for MEG and 1921 for EEG. To ensure voxel-wise correspondence across modalities, the four MEG voxels that could not be reconstructed in EEG were excluded from the MEG vector prior to analysis.

Normality of the natural frequency distributions was assessed using the Kolmogorov–Smirnov test, which revealed that none of the maps followed a Gaussian distribution (all *p* < .001). Therefore, non-parametric Spearman rank correlations were used to quantify the similarity between maps.

Because the eyes-open condition was the only one available across all recording modalities, correlation analyses were restricted to this condition and conducted in two steps. First, we assessed the correspondence between MEG (MEG-EO) and high-density EEG (EEG-64-EO). Second, we evaluated the correspondence between high-and low-density EEG conditions (EEG-64-EO and EEG-32-EO).

### 2.6. Differences in natural frequencies across recording modalities and eye conditions

To examine condition-specific changes in natural frequencies, we performed voxel-wise comparisons of natural frequency maps across recording modalities and eye conditions. Comparisons across recording modalities were conducted between MEG-EO and EEG-64-EO, as well as between EEG-64-EO and EEG-32-EO. Comparisons between eyes-open and eyes-closed conditions were performed only for the 64-channel EEG configuration (i.e., EEG-64-EO vs. EEG-64-EC). Analyses were based on the previously computed single-subject maps, yielding one frequency value per voxel per participant. As described above, only the 1921 voxels common to EEG and MEG were included to ensure voxel-wise correspondence across modalities.

Because individual voxels can show substantial variability in frequency across participants (e.g., 5 vs. 25 Hz in frontal voxels), conventional summary statistics based on measures of central tendency are not appropriate. To address this limitation, and following Herrera-Morueco et al. (2026), we computed, for each voxel, participant, and condition, the proportion of neighboring voxels (within a 1.5 cm radius) exhibiting each frequency. This approach yields normalized local frequency distributions that capture the prevalence of each frequency within the spatial neighborhood of each voxel.

Condition differences were then assessed using a non-parametric permutation framework combined with effect size estimation (Herrera-Morueco et al., 2026). For each voxel and comparison, Cohen’s *d* was computed for each frequency of the normalized local frequency distributions, and the maximum *d* value was retained as the test statistic. Independent-samples Cohen’s *d* was used for MEG vs. EEG comparisons (given that participants differed between modalities), whereas repeated-measures Cohen’s *d* was applied when comparing EEG density configurations and eye conditions (as the same participants were included). Statistical significance was assessed by generating a null distribution under the hypothesis of no condition differences through 1000 label permutations. For each permutation, condition labels were randomly shuffled, and the maximum statistic across all voxels was stored to correct for multiple comparisons. Voxels were considered statistically significant when observed *d* values exceeded the 95th percentile of the null distribution.

For visualization purposes, we selected a subset of voxels exhibiting statistically significant effects, focusing on local maxima in the statistical maps and their spatially symmetric counterparts.

## 3. Results

### 3.1. Moderate similarity between high-density EEG and MEG natural frequency maps, and strong similarity between high- and low-density EEG

To assess the similarity of natural frequency maps across recording modalities and EEG densities, we computed pairwise Spearman rank correlations between the group-level maps obtained during eyes-open conditions from MEG (MEG-EO), high-density EEG (EEG-64-EO), and low-density EEG (EEG-32-EO). Visual inspection of the maps (Figure 1) reveals a highly similar spatial organization of natural frequencies across modalities, with consistent anterior-to-posterior and medial-to-lateral gradients of increasing frequency.

**Figure 1.**
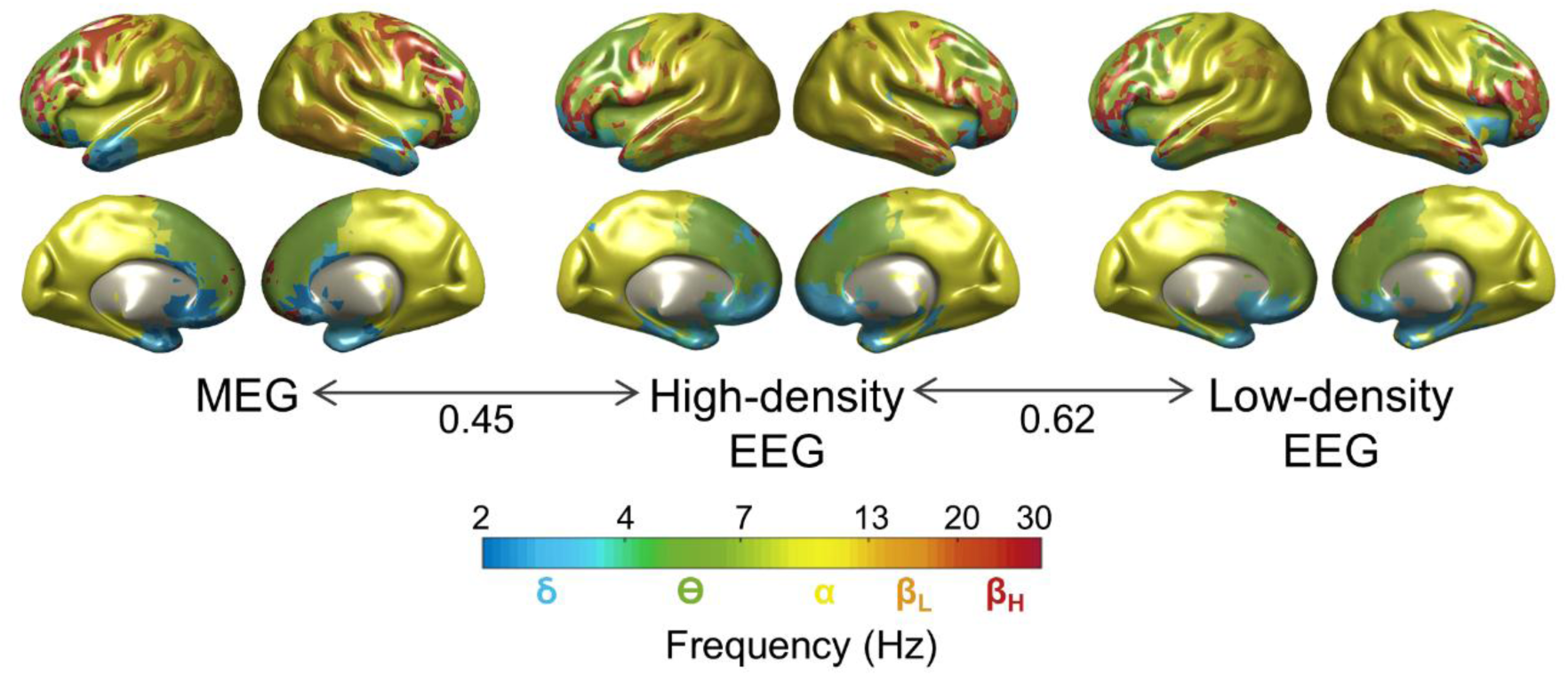
Natural frequencies across recording modalities. From left to right, correlation between MEG (MEG-EO) and high-density EEG (EEG-64-EO), and between high- (EEG-64-EO) and low-density EEG (EEG-32-EO). The color code indicates different oscillatory frequencies, ranging from the delta to the high-beta band.

Quantitatively, a moderate correlation was observed between MEG and high-density EEG maps (*ρ* = 0.45), indicating that both modalities capture broadly similar spatial patterns of natural frequencies at the group level. A visual comparison of the maps suggested that differences between MEG and EEG were most pronounced in lateral frontal and temporal regions, whereas posterior and medial areas showed highly similar frequency patterns.

The comparison between high- and low-density EEG showed a stronger correlation (*ρ* = 0.62), and the maps appeared nearly indistinguishable across most regions, suggesting that reducing the number of EEG channels has a limited impact on the resulting natural frequency brain maps.

These results indicate that, while the overall spatial pattern of natural frequencies is preserved across modalities and EEG densities, regional differences—particularly in lateral frontal and temporal areas—contribute to the moderate correspondence observed between MEG and EEG.

### 3.2. Frontal regions show the largest differences between MEG and EEG, while natural frequencies remain consistent across EEG densities

To identify the cortical regions underlying the observed differences between recording modalities under eyes-open conditions, we performed voxel-wise statistical comparisons of natural frequency distributions. Figure 2a shows effect size maps (Cohen’s *d*) for the comparison between MEG and high-density EEG. Significant differences were primarily observed in frontal regions, with the largest effects located in the frontolateral cortex, followed by frontomedial and cingulate areas, as well as in sensorimotor areas and temporal cortex. In contrast, posterior regions did not show significant differences between modalities.

**Figure 2.**
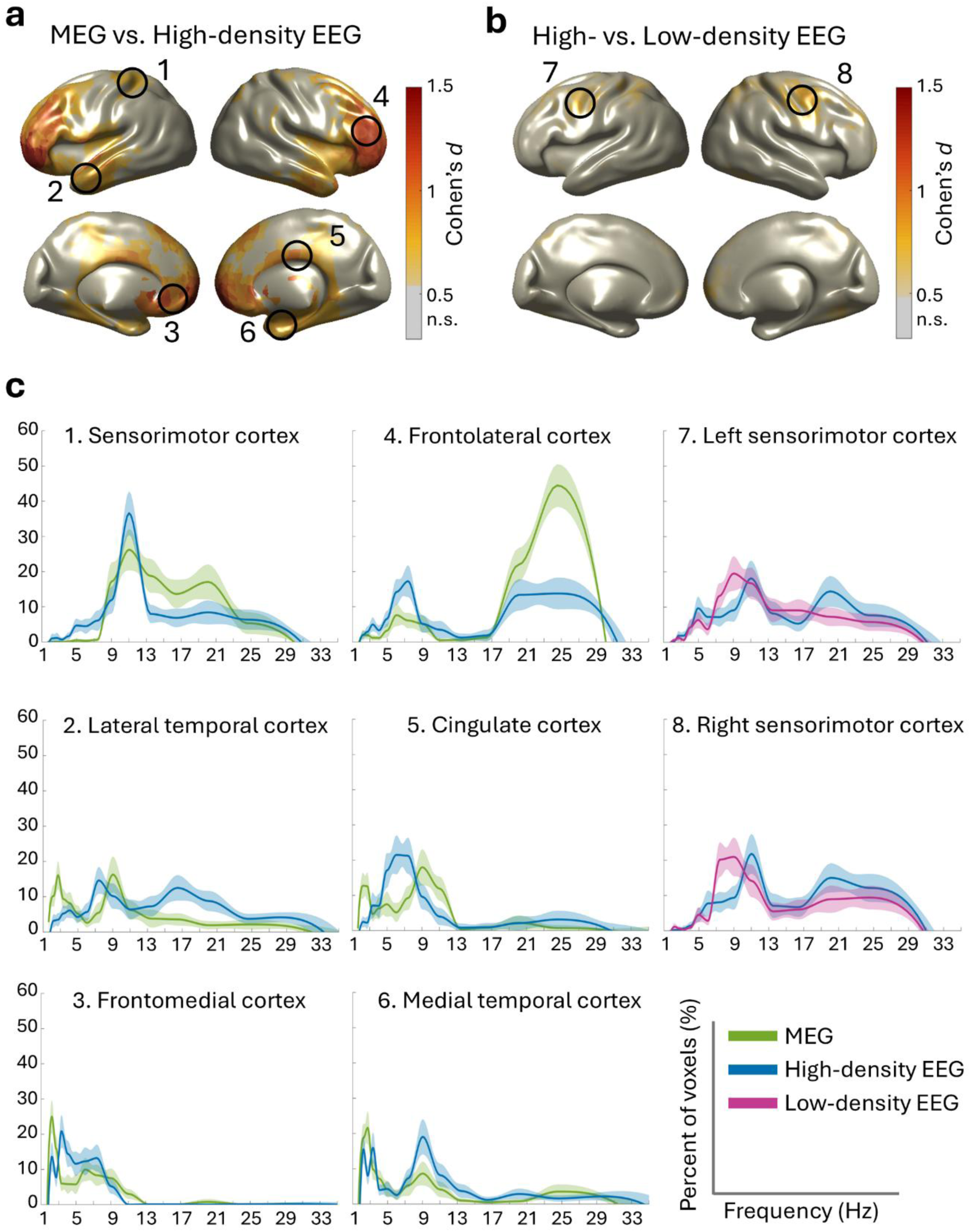
Differences in natural frequencies across recording modalities and EEG densities. (**a**) Voxel-wise effect size map (Cohen’s *d*) for the MEG (MEG-EO) vs. high-density EEG (EEG-64-EO) comparison. (**b**) Voxel-wise effect size map for the high- (EEG-64-EO) vs. low-density (EEG-32-EO) EEG comparison. Significant regions are shown using an orange-to-red color scale; n.s., not significant. Black circles indicate the location of significant voxels displayed in panel (c). (**c**) Local frequency distributions of representative significant voxels, indicating the prevalence of natural frequencies within their spatial neighborhood. MEG is depicted in green, high-density EEG in blue, and low-density EEG in pink. Shaded areas represent the standard error of the mean (SEM).

The regional frequency distributions illustrated in Figure 2c further clarify these effects by showing the percentage of voxels oscillating at each frequency within a local neighborhood. In frontolateral regions, MEG showed a predominance of voxels exhibiting higher frequencies compared to high-density EEG (25 vs. 7 Hz). In medial frontal and temporal regions, as well as in the lateral temporal cortex, MEG exhibited a subtle shift towards lower frequencies relative to EEG. In the cingulate cortex, MEG showed a bimodal pattern, with some voxels oscillating in the delta range (∼2 Hz), and others in the alpha range (∼9 Hz), whereas the majority of voxels in EEG presented frequencies in the intermediate theta range (∼7 Hz). Finally, over sensorimotor regions, most voxels showed a clear peak in the alpha range (∼10 Hz) in EEG, whereas in MEG this pattern was more broadly distributed across alpha and beta frequencies.

The comparison between high- and low-density EEG (Figure 2b) revealed very limited significant effects, indicating that reducing the number of EEG channels does not produce major changes in the spatial distribution of natural frequencies. This is further supported by the nearly overlapping frequency distributions observed in sensorimotor and premotor regions for both EEG densities, with a predominance of natural frequencies in the alpha and beta ranges (Figure 2c).

Together, these results indicate that the main differences in natural frequencies between MEG and EEG were driven by frontal and temporal regions. Despite these differences, frequency distributions across regions remained largely comparable between modalities, showing similar overall profiles and peak frequency values, particularly in posterior cortices. Furthermore, natural frequencies remained consistent across EEG densities.

### 3.3. Posterior regions show the largest differences during eye closure

To characterize the effects of eye closure on the spatial organization of natural frequencies, we first examined the group-level natural frequency map obtained during the eyes-closed condition, followed by voxel-wise statistical comparisons between the eyes-closed (EEG-64-EC) and eyes-open (EEG-64-EO) conditions.

Figure 3a shows the group-level natural frequency map for the eyes-closed condition. Qualitatively, the map revealed a clear predominance of alpha-band frequencies in posterior cortices, while frontal areas showed more heterogeneous frequency patterns spanning primarily the delta, theta, and alpha ranges.

**Figure 3.**
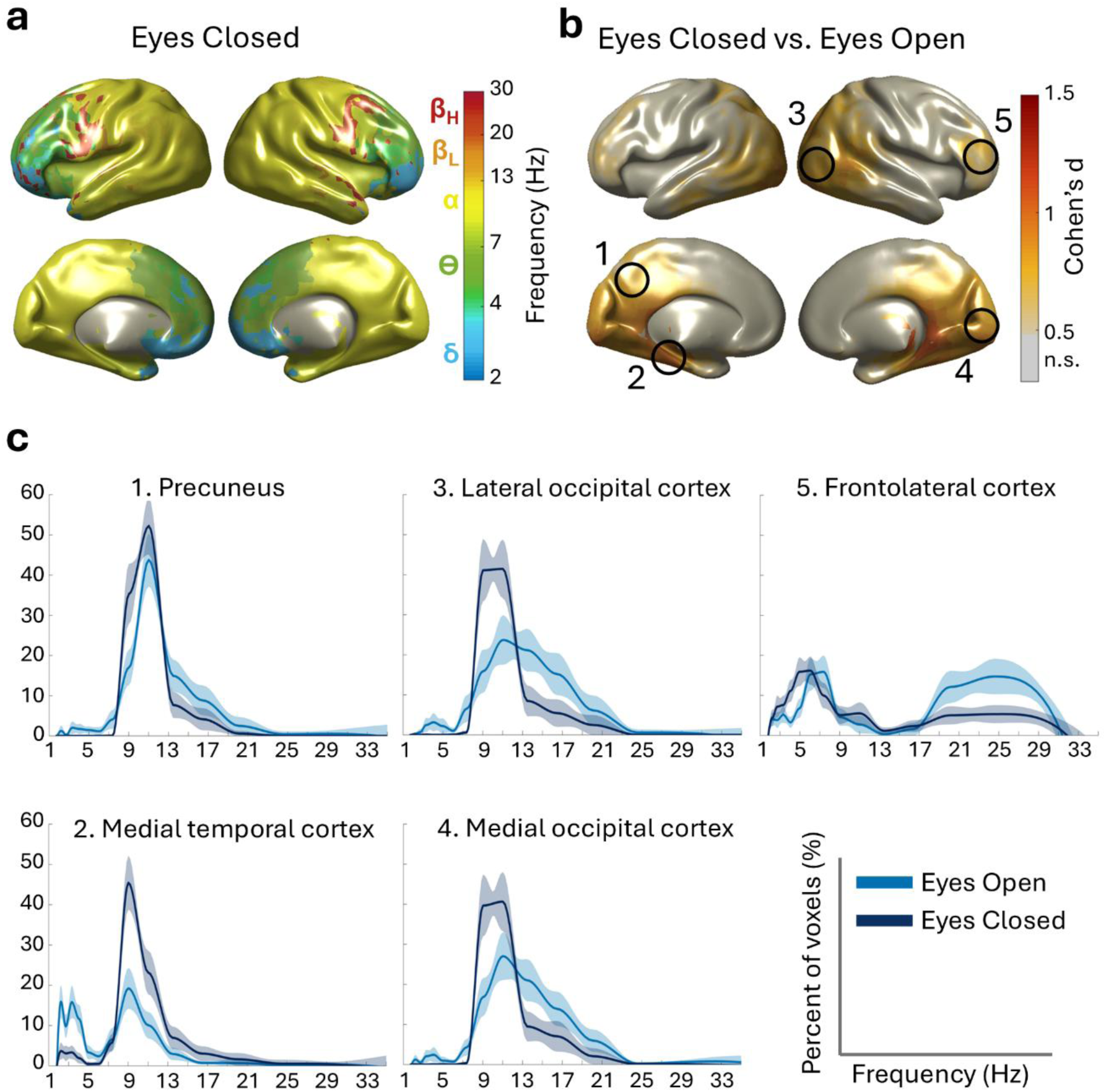
Differences in natural frequencies between eyes-open and eyes-closed conditions. (**a**) Group-level natural frequency map for the eyes-closed condition using EEG (EEG-64-EC). (**b**) Voxel-wise effect size map (Cohen’s *d*) for the eyes-closed (EEG-64-EC) vs. eyes-open (EEG-64-EO) comparison. Significant regions are shown using an orange-to-red color scale; n.s., not significant. Black circles indicate the location of significant voxels displayed in panel (c). (**c**) Local frequency distributions of representative significant voxels, indicating the prevalence of natural frequencies within their spatial neighborhood. Eyes open is depicted in light blue and eyes closed in dark blue. Shaded areas represent the standard error of the mean (SEM).

Figure 3b displays the voxel-wise effect size map (Cohen’s *d*) for the comparison between eyes-closed and eyes-open EEG. Significant differences were primarily observed in posterior regions, including medial and lateral occipital cortices and the precuneus, with smaller effects in frontolateral and medial temporal regions.

Figure 3c illustrates the percentage of voxels oscillating at each frequency for each region. In the precuneus, both conditions showed a clear predominance of alpha activity, although slightly stronger in the eyes-closed condition. In the medial temporal cortex, eyes-closed EEG presented a clear predominance of alpha frequencies, whereas eyes-open EEG showed a bimodal pattern with peaks in the alpha (∼9 Hz) and delta (∼3 Hz) ranges. In both the lateral and medial occipital cortices, the eyes-closed condition exhibited a sharp alpha peak (∼10 Hz), whereas eyes-open EEG showed a broader distribution extending toward higher frequencies. Similarly, in the frontolateral cortex, the eyes-open condition showed a wider distribution of frequencies relative to the eyes-closed condition, extending into the beta range.

### 3.4. Stability of single-subject natural frequency patterns across EEG conditions

Figure 4 shows examples of single-subject natural frequency maps for the three EEG conditions: eyes open with high-density EEG (EEG-64-EO), eyes open with low-density EEG (EEG-32-EO), and eyes closed with high-density EEG (EEG-64-EC).

**Figure 4.**
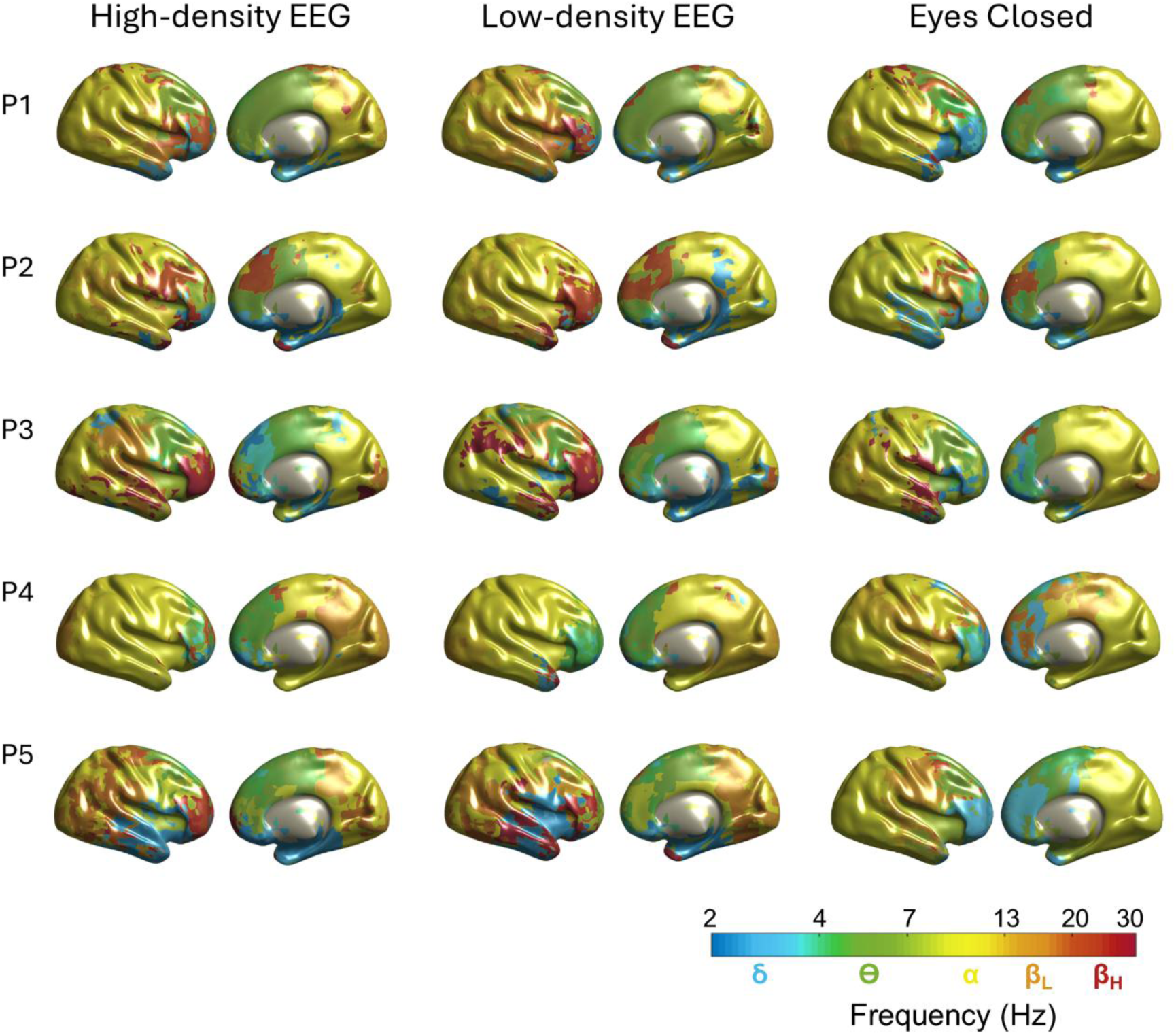
Single-subject maps of natural frequencies across EEG conditions. Examples from five participants (P) are shown in rows, with each column representing one condition: eyes open with high-density EEG (EEG-64-EO), eyes open with low-density EEG (EEG-32-EO), and eyes closed with high-density EEG (EEG-64-EC). For simplicity, only lateral views of the right hemisphere and medial views of the left hemisphere are displayed. The color code indicates different oscillatory frequencies, ranging from the delta to the high-beta band.

A qualitative comparison between high- and low-density EEG maps under eyes-open conditions shows that each participant exhibits a characteristic pattern of natural frequencies that remains stable across EEG density conditions. When comparing the eyes-closed to the eyes-open conditions, a shift toward lower frequencies can be observed, consistent with the results reported in the previous section.

## 4. Discussion

In this study, we transferred the MEG-based methodology for mapping brain natural frequencies (Capilla et al., 2022) to EEG and evaluated its robustness across recording modalities, EEG channel densities, and physiological states (eyes open and eyes closed). Our results showed that EEG was able to recover a spatial organization of natural frequencies comparable to that obtained with MEG, that this organization was largely preserved when reducing EEG density from 64 to 32 channels, and that the method was sensitive to well-known physiological modulations such as eye closure.

The comparison between MEG and high-density EEG maps revealed a moderate but clear similarity in the organization of natural frequencies across the cortex. Both recording modalities exhibited comparable posterior-to-anterior and medial-to-lateral gradients of increasing frequency. This overall pattern is consistent with previous reports of spatial spectral organization in MEG (Capilla et al., 2022; Keitel & Gross, 2016; Lew et al., 2021; Mahjoory et al., 2020; Ramkumar et al., 2014) and EEG (Chen et al., 2008; Congedo et al., 2010; Janiukstyte et al., 2023). Our results extend these findings by showing that this spatial pattern can be recovered without relying on predefined frequency bands, reinforcing the idea that these gradients reflect genuine cortical oscillatory properties.

This result gains relevance in light of the well-known differences between the two modalities in terms of signal properties. EEG and MEG differ in their sensitivity to source orientation, depth, and size, as well as in the distortion of electric—but not magnetic—field propagation by head tissues (Bénar et al., 2019; Goldenholz et al., 2009; Klamer et al., 2015; Malmivuo, 2012; Srinivasan et al., 2006). In this context, the main voxel-wise differences observed in frontal, sensorimotor, and temporal regions can be interpreted in terms of the spatial filtering properties and differential sensitivity of EEG and MEG to cortical source configurations of varying size and orientation. Scalp EEG preferentially captures radial components of large, synchronous cortical sources located on gyral crowns, whereas MEG is more sensitive to tangential components arising from sources located along sulcal walls and at the boundaries of distributed cortical activity patterns (Bénar et al., 2019; Goldenholz et al., 2009; Malmivuo, 2012; Srinivasan et al., 2006).

This framework provides a parsimonious explanation for modality-specific differences in the relative prominence of frequency components. For example, activity dominated by lower-frequency rhythms reflecting broadly synchronized cortical populations (e.g., delta, theta, alpha) tends to be more salient in EEG, whereas higher-frequency components arising from more spatially localized activity may be more readily detected by MEG, owing to its reduced dependence on skull conductivity effects and its finer effective spatial resolution (Bénar et al., 2019; Malmivuo, 2012; Srinivasan et al., 2006)—note that MEG’s spatial resolution is not inherently superior to EEG when using comparable channel densities (Klamer et al., 2015; Liu et al., 2002). In regions where large, coherent radial sources coexist with smaller, more local tangential sources, such as posterior cortices (Srinivasan et al., 2006), spectral profiles appear highly similar across EEG and MEG. Consistent with this interpretation, Janiukstyte et al. (2023) reported moderate-to-strong spatial correlations between EEG and MEG across canonical frequency bands, with delta and alpha showing the highest concordance, while correlations in higher-frequency bands, such as beta, were weaker.

Overall, our findings suggest that both EEG and MEG capture complementary aspects of the same underlying cortical organization, and that the regional discrepancies reflect genuine differences in the spatial scales of cortical synchronization rather than modality-specific artifacts. However, additional methodological factors may also contribute to these effects. Differences in source reconstruction procedures, such as the use of a standard MRI template for EEG versus individual anatomical data for MEG, may introduce regional variability (Klamer et al., 2015). Moreover, the transmission of electric—but not magnetic—signals from cortex to scalp is frequency-dependent: higher-frequency oscillations are more strongly attenuated due to partial cancellation of activity from neighboring cortical sources, whereas lower-frequency rhythms are less affected and therefore more reliably detected at the scalp (Bénar et al., 2019; Malmivuo, 2012; Pfurtscheller & Cooper, 1975).

An additional factor that may account for the observed differences between MEG and EEG—absent in the comparison between EEG systems of different densities—is that the participants were not the same across modalities. Notably, the largest discrepancies between MEG and EEG were found in frontal regions, which have been shown to contribute strongly to individual identification from natural frequency maps, likely due to their higher inter-individual variability (Arana et al., 2025). Thus, an optimal design would involve recordings from the same participants using both EEG and MEG, ideally acquired simultaneously.

Comparisons between high- and low-density EEG further suggest that variability in natural frequency mapping is not primarily driven by channel sampling density. The strong similarity observed across both EEG configurations—at the group level and in single-subject maps—together with the negligible differences identified in voxel-wise analyses, indicates that the spatial organization of natural frequencies does not critically depend on high channel density. Individual participants exhibited highly stable patterns across densities, and voxel-wise statistical comparisons revealed only minor discrepancies in sensorimotor cortices, likely reflecting the coexistence of alpha and beta rhythms in these regions (Salmelin & Hari, 1994; Babiloni et al., 2016). These findings challenge the common assumption that high-density EEG is required for reliable source-level spatial characterization. Instead, they suggest that the organization captured by natural frequency mapping reflects large-scale cortical gradients that can be robustly reconstructed even with reduced sensor coverage. From a methodological perspective, this robustness is likely related to the fact that the approach does not rely on fine-grained spatial details, but rather on consistent spectral patterns across neighboring voxels (Arana et al., 2025). From a practical standpoint, these findings have direct implications for clinical applicability, where low-density EEG systems are much more commonly used.

The eyes-closed condition revealed a pronounced predominance of natural frequencies in the alpha range (8–13 Hz) at both the group and single-subject levels. Voxel-wise statistics indicated that the main differences between the eyes-open and eyes-closed conditions were located in posterior regions, including both medial and lateral occipital cortices, while more subtle differences were observed in frontolateral and medial temporal areas.

The posterior pattern directly corresponds to one of the most robust and well-established phenomena in human electrophysiology: posterior alpha enhancement during eye closure (Berger, 1929; see Quigley, 2022 for a review). Numerous studies using either EEG or MEG have demonstrated this effect when comparing eyes-open and eyes-closed conditions (Barry et al., 2007; Chen et al., 2008; Geller et al., 2014; Petro et al., 2022). Moreover, the shift toward lower frequencies observed in frontal regions during eye closure is consistent with previous findings in spectral power (Chen et al., 2008; Petro et al., 2022), along with the decrease in power at higher frequencies (Barry et al., 2007; Geller et al., 2014). It has been proposed that frontal beta activity is associated with higher-level processing and integration of visual information (Di Dona & Ronconi, 2023), suggesting that its reduction during eye closure may be linked to reduced visual processing demands. In medial temporal regions, the eyes-closed condition showed a clear dominance of voxels presenting only alpha frequencies, in contrast to the bimodal delta–alpha distribution observed during the eyes-open condition. This finding aligns with Barry et al. (2007), who reported increased alpha power not only over the occipital cortex but also extending to other regions such as the temporal cortex. Thus, the predominance of alpha activity likely reflects global resting-state arousal, whereas changes in other frequency bands appear to reflect localized cortical processing of visual input rather than a mere increase in arousal (Barry et al., 2007). These results show that natural frequency mapping does not merely capture a static spatial gradient, but rather a physiologically meaningful cortical organization that dynamically reflects changes in brain state.

Taken together, our findings show that EEG, including low-density configurations, can reliably map natural cortical frequencies and recover the cortical gradients previously observed with MEG. Because EEG is widely available in clinical and hospital settings, this methodology becomes readily transferable to contexts where MEG is impractical. This opens the possibility of using EEG-based natural frequency mapping as a non-invasive tool for studying brain oscillations and identifying alterations in neuropsychiatric conditions, including so-called oscillopathies, as well as for tracking longitudinal changes in both healthy and clinical populations.

## 5. Conclusion

Natural frequencies may be regarded as an intrinsic property of cortical organization that can be reliably recovered with EEG, even at low channel densities and across different physiological states. Although EEG and MEG differ in their inherent sensitivity to neural source configuration, these differences do not undermine the validity of EEG-based natural frequency mapping. Together, these findings demonstrate that the concept of natural frequencies is not modality-specific and can be extended beyond specialized MEG research environments. This opens the possibility of using EEG-based natural frequency mapping as a practical, non-invasive tool for studying brain oscillations and their alterations in neuropsychiatric conditions.

## Data availability

The EEG dataset used in this study is available at OpenNeuro: Resting-state EEG data before and after cognitive activity across the adult lifespan and a 5-year follow-up (doi: 10.18112/openneuro.ds005385.v1.0.2; Getzmann et al., 2024). MEG data are available at https://www.mcgill.ca/bic/neuroinformatics/omega from OMEGA: The Open MEG Archive (Niso et al., 2016).

## Code availability

The DISCOVER-EEG pipeline used for EEG-data preprocessing pipeline is available at https://github.com/crisglav/discover-eeg (Gil Ávila et al., 2023). The code for MEG and EEG preprocessing and all subsequent analyses is available at https://github.com/necog-UAM.

## Acknowledgments

We are thankful to all the researchers and technicians involved in the Dortmund Vital Study for recording and making publicly available the EEG dataset, as well as to all the researchers and technicians involved in OMEGA: The Open MEG Archive. The authors also thank Jorge San-Segundo and Joachim Gross for their valuable advice regarding our research.

This work was supported by Ministerio de Ciencia, Innovación y Universidades / Agencia Estatal de Investigación, Spain / FEDER/FSE+, UE (MCIU/AEI/10.13039/501100011033 /FEDER/FSE+, UE; PID2021-125841NB-I00 and PID2024-161032NB-I00 to AC, PRE2022-101613 to ES, and JDC2023-050403-I to MM); and the Comunidad de Madrid, Spain (IND2022/SOC-23652 to LA and PIPF-2024/SAL-GL-34549 to JJHM).

## Author contributions

Lydia Arana: Conceptualization, Data curation, Formal analysis, Methodology, Software, Visualization, Writing – original draft preparation, Writing – review & editing; Juan José Herrera-Morueco: Methodology, Software, Writing – review & editing; María Melcón: Data curation, Software, Writing – review & editing; Enrique Stern: Resources, Software, Writing – review & editing; Sandra Pusil: Conceptualization, Writing – review & editing, Supervision; Almudena Capilla: Conceptualization, Funding acquisition, Methodology, Resources, Writing – original draft preparation, Writing – review & editing, Supervision.

## Supplementary material

### MEG-EO k-means centroids

**Supplementary Figure 1.**
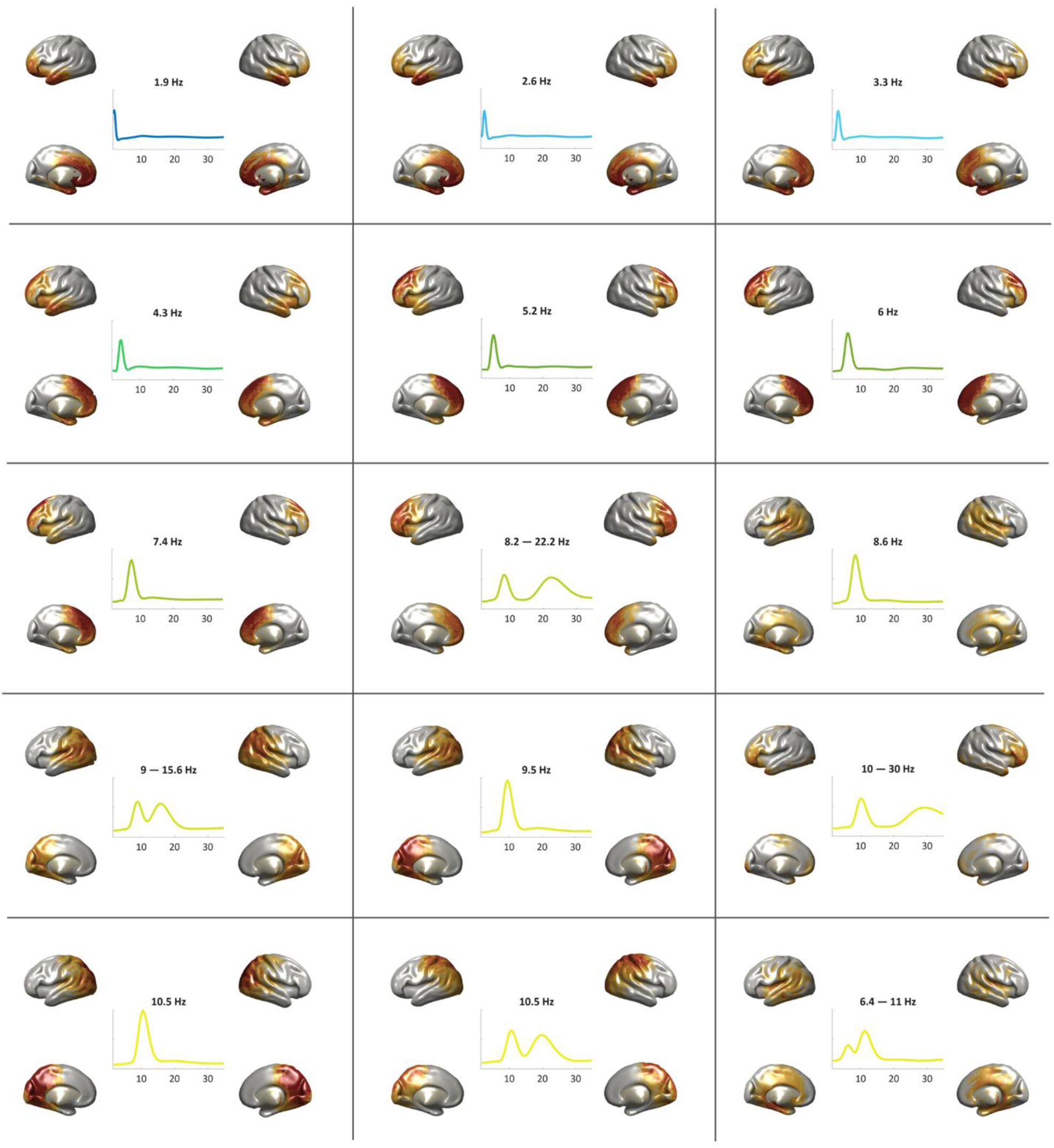
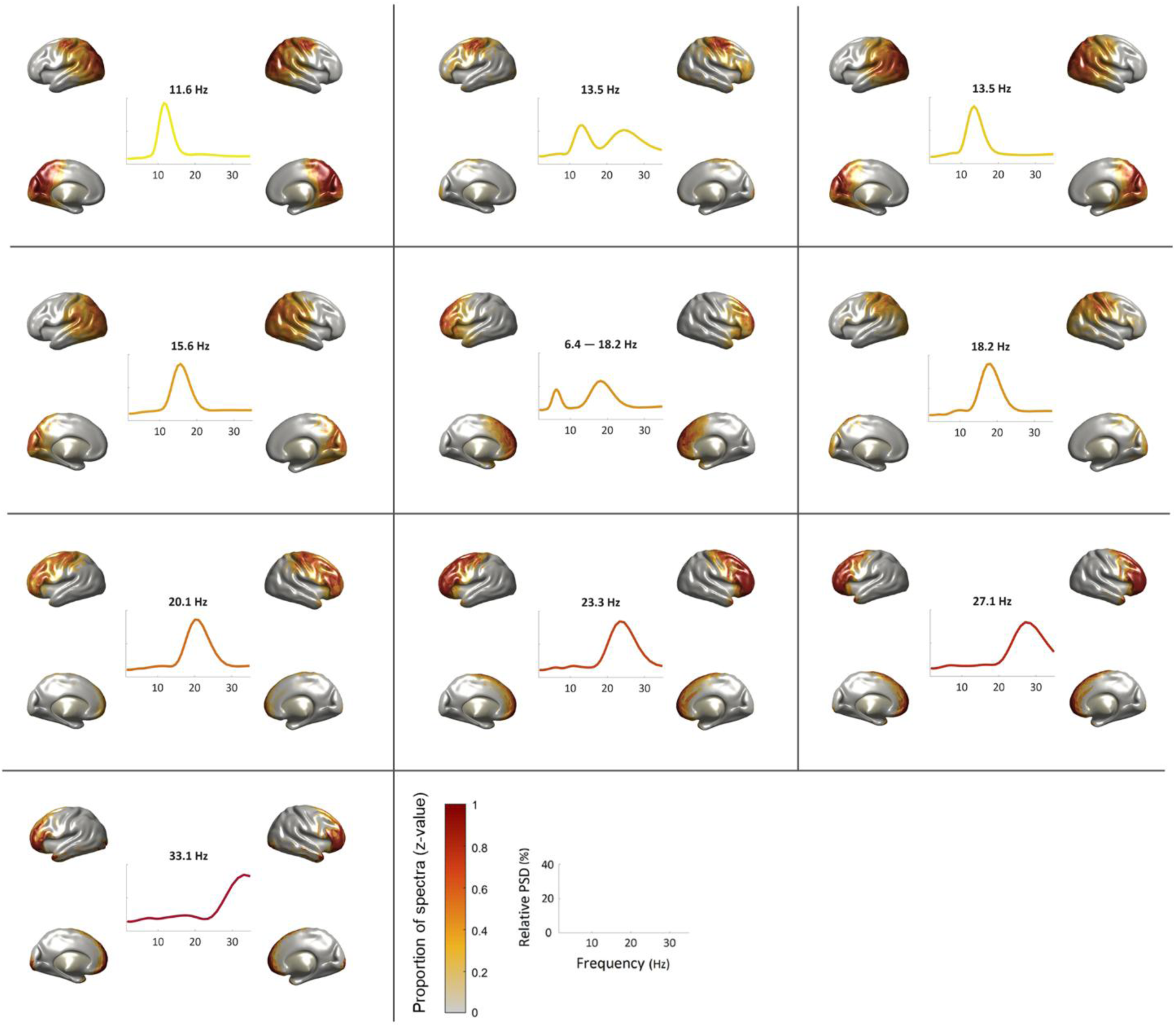
Brain generators and centroid power spectra for the MEG-EO (MEG eyes-open) k-means clustering. The figure shows the 25 cluster centroids resulting from the whole-group clustering. Each panel presents the centroid power spectra, peak frequency, and spatial distribution, i.e., the proportion of spectra assigned to that cluster normalized across voxels (z-score). In cases where clusters exhibited two peaks and one was identified as a harmonic of the other, only the fundamental frequency was considered.

### EEG-64-EO k-means centroids

**Supplementary Figure 2.**
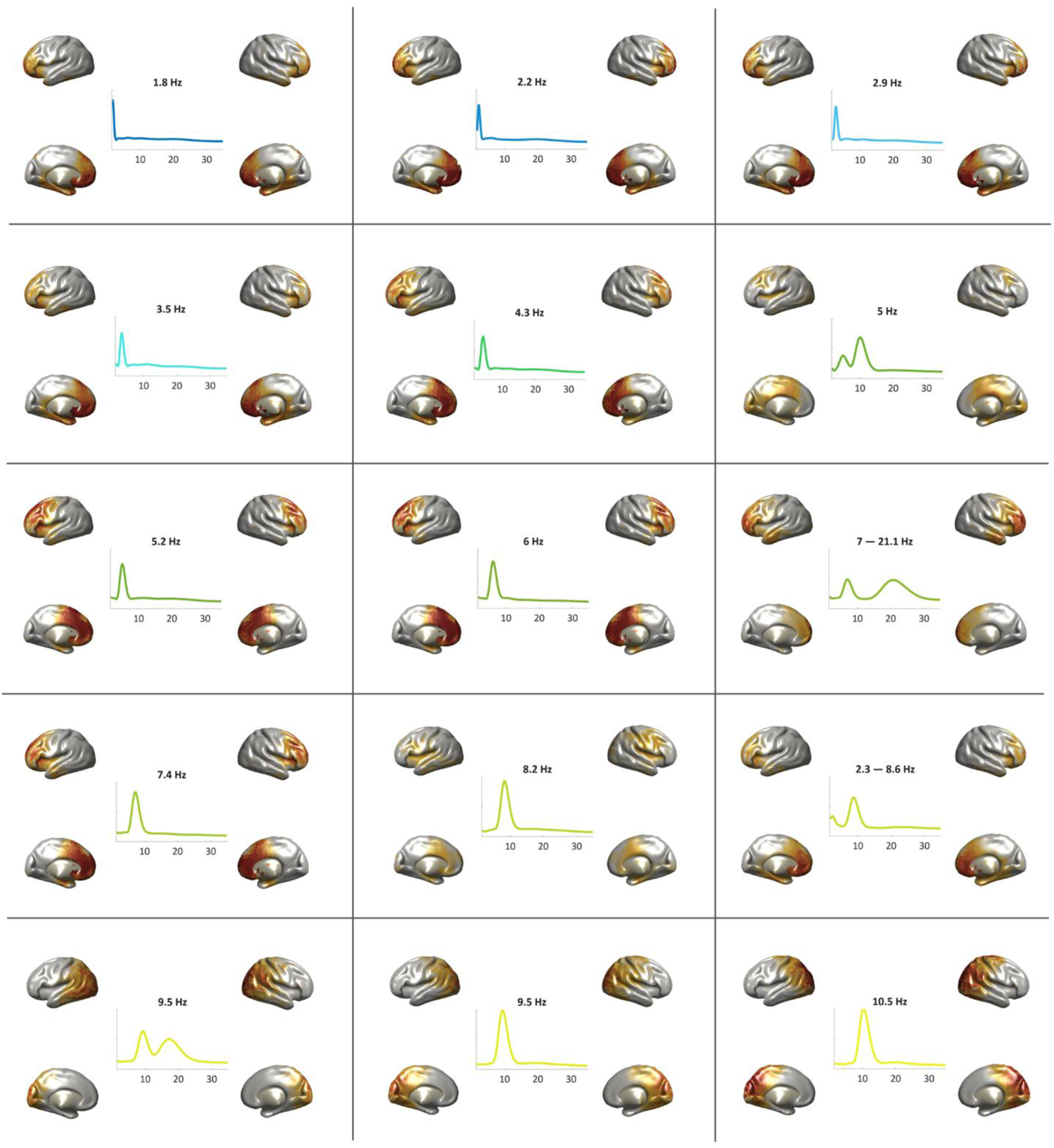
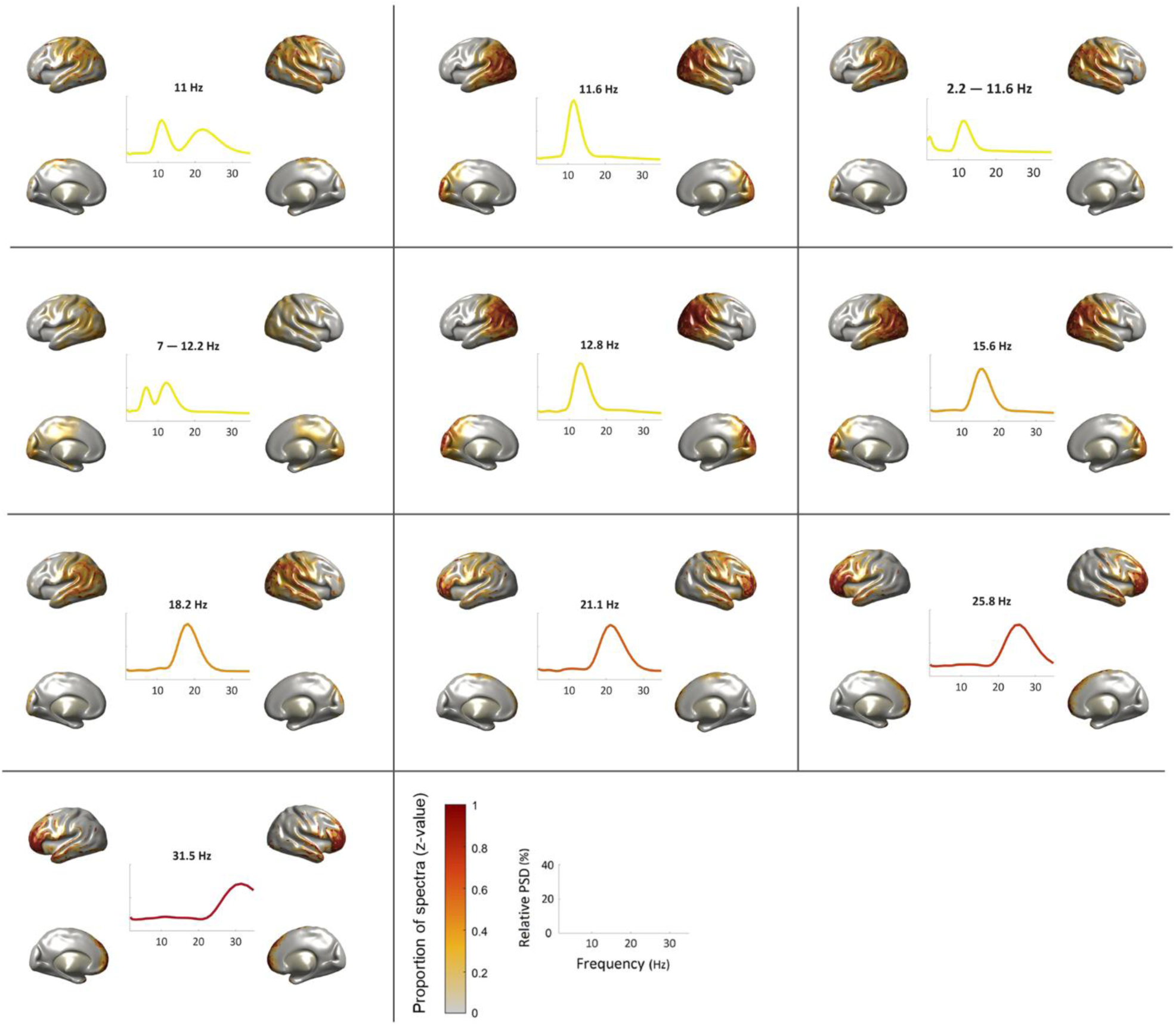
Brain generators and centroid power spectra for the EEG-64-EO (EEG 64-channels eyes-open) k-means clustering. The figure shows the 25 cluster centroids resulting from the whole-group clustering. Each panel presents the centroid power spectra, peak frequency, and spatial distribution of each cluster, i.e., the proportion of spectra assigned to that cluster normalized across voxels (z-score). In cases where clusters exhibited two peaks and one was identified as a harmonic of the other, only the fundamental frequency was considered.

### EEG-32-EO k-means centroids

**Supplementary Figure 3.**
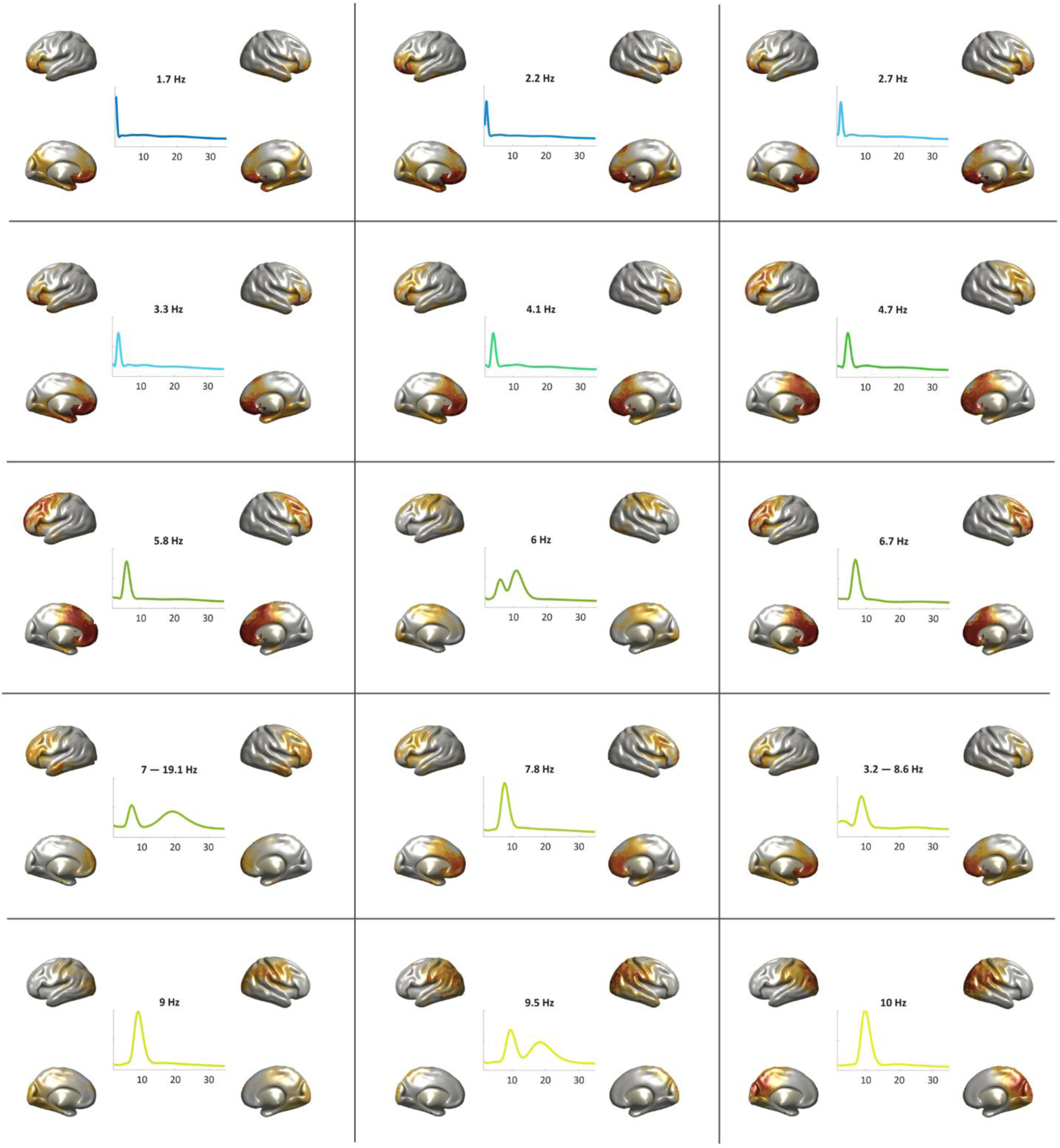
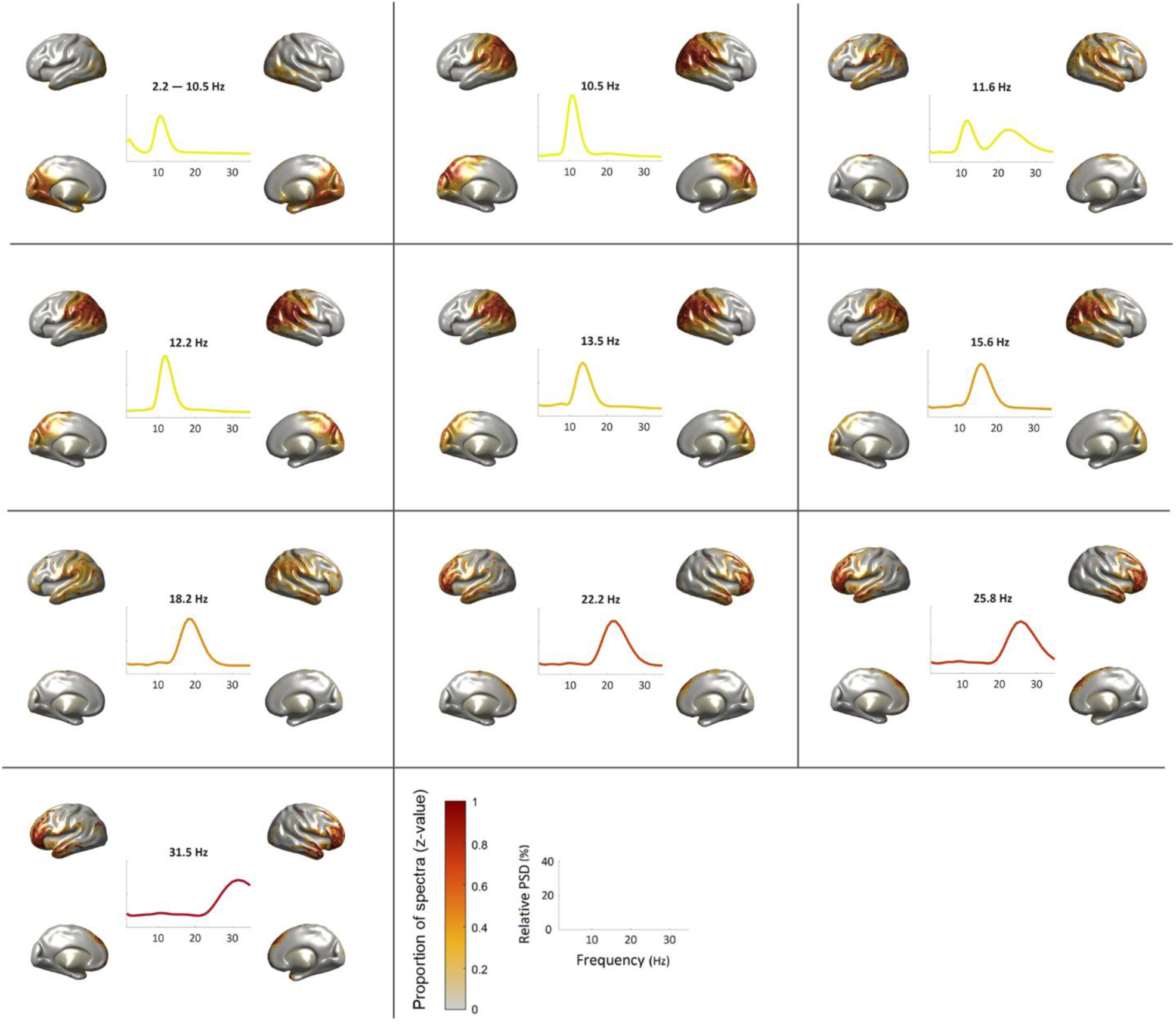
Brain generators and centroid power spectra for the EEG-32-EO (EEG 32-channels eyes-open) k-means clustering. The figure shows the 25 cluster centroids resulting from the whole-group clustering. Each panel presents the centroid power spectra, peak frequency, and spatial distribution of each cluster, i.e., the proportion of spectra assigned to that cluster normalized across voxels (z-score). In cases where clusters exhibited two peaks and one was identified as a harmonic of the other, only the fundamental frequency was considered.

### EEG-64-EC k-means centroids

**Supplementary Figure 4.**
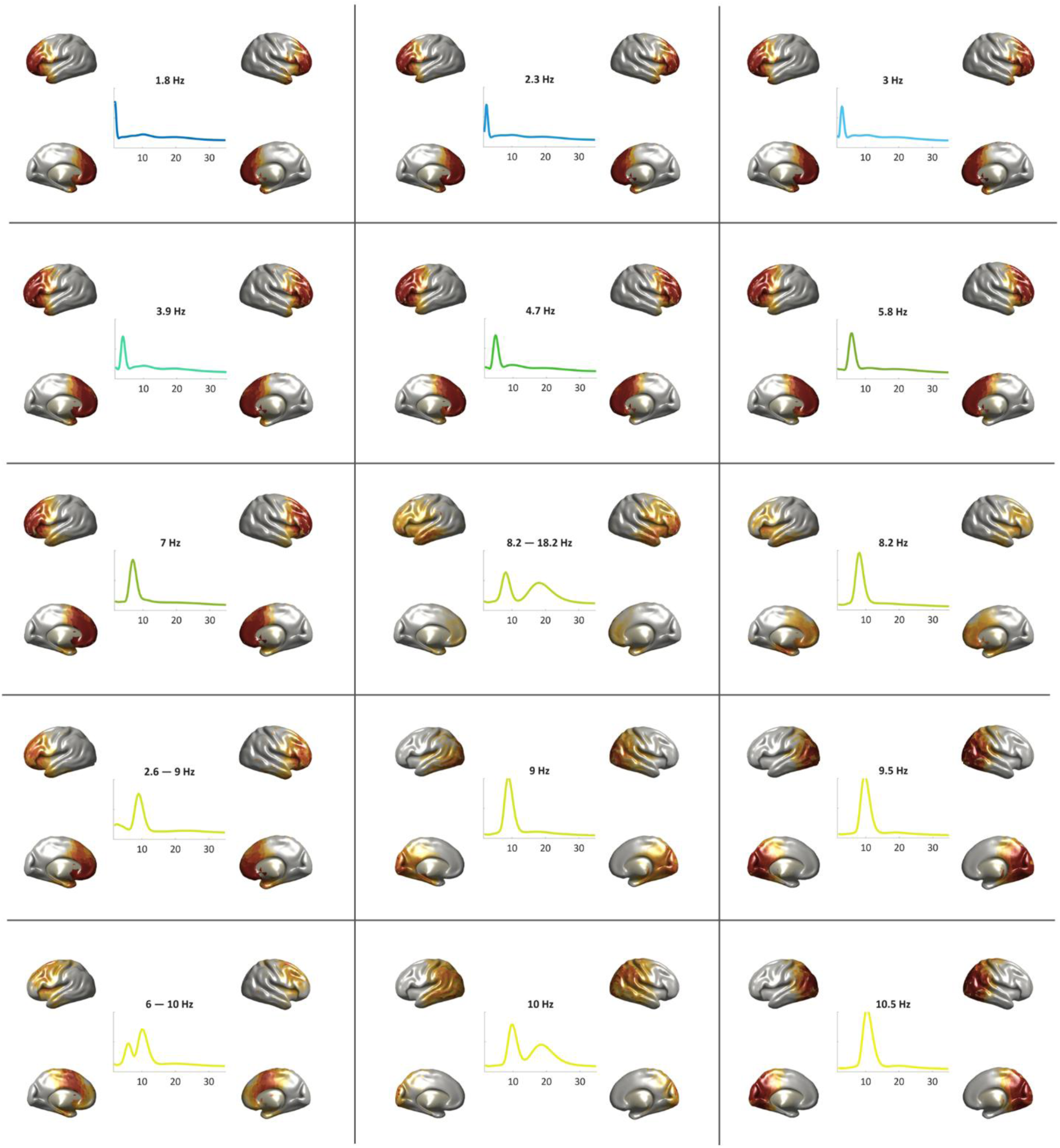
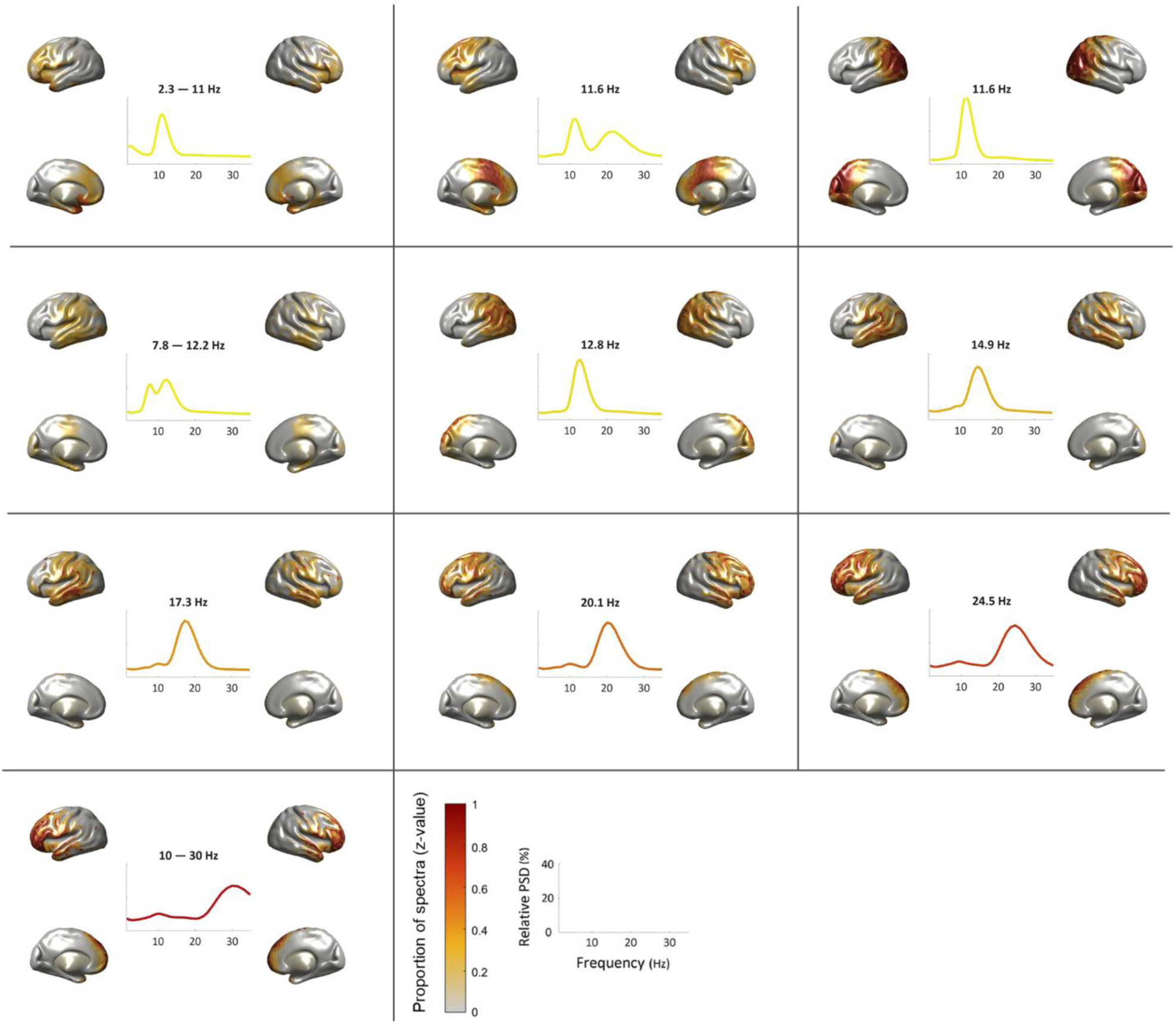
Brain generators and centroid power spectra for the EEG-64-EC (EEG 64-channel eyes-closed) k-means clustering. The figure shows the 25 cluster centroids resulting from the whole-group clustering. Each panel presents the centroid power spectra, peak frequency, and spatial distribution of each cluster, i.e., the proportion of spectra assigned to that cluster normalized across voxels (z-score). In cases where clusters exhibited two peaks and one was identified as a harmonic of the other, only the fundamental frequency was considered.

## References

1. Amengual, J. L., Stengel, C., Moreau, T., Adam, C., Chavez, M., & Valero-Cabré, A. (2019). Perturbation-based mapping of natural frequencies with direct intracranial stimulation of the human brain. BioRxiv. 10.1101/718064

2. Arana, L., Herrera-Morueco, J. J., Santonja, J., & Capilla, A. (2025). Identifying individuals from their brain natural frequency fingerprints. Scientific Reports, 15(1), 22492. 10.1038/s41598-025-05632-7

3. Babiloni, C., Del Percio, C., Vecchio, F., Sebastiano, F., Di Gennaro, G., Quarato, P. P., Morace, R., Pavone, L., Soricelli, A., Noce, G., Esposito, V., Rossini, P. M., Gallese, V., & Mirabella, G. (2016). Alpha, beta and gamma electrocorticographic rhythms in somatosensory, motor, premotor and prefrontal cortical areas differ in movement execution and observation in humans. Clinical Neurophysiology, 127(1), 641–654. 10.1016/j.clinph.2015.04.068

4. Barry, R. J., Clarke, A. R., Johnstone, S. J., Magee, C. A., & Rushby, J. A. (2007). EEG differences between eyes-closed and eyes-open resting conditions. Clinical Neurophysiology, 118(12), 2765–2773. 10.1016/j.clinph.2007.07.028

5. Bénar, C. G., Grova, C., Jirsa, V. K., & Lina, J. M. (2019). Differences in MEG and EEG power-law scaling explained by a coupling between spatial coherence and frequency: A simulation study. Journal of Computational Neuroscience, 47(1), 31–41. 10.1007/s10827-019-00721-9

6. Berger, H. (1929). Uber das Elektrenkephalogramm des Menschen. Archiv Für Psychiatrie Und Nervenkrankheiten, 87, 527–570.

7. Besl, P., & McKay, N. (1992). A method for registration of 3D shapes. IEEE Transactions on Pattern Analysis and Machine Intelligence, 14(2), 239–256.

8. Boersma, M., Smit, D. J. A., De Bie, H. M. A., Van Baal, G. C. M., Boomsma, D. I., De Geus, E. J. C., Delemarre-van De Waal, H. A., & Stam, C. J. (2011). Network analysis of resting state EEG in the developing young brain: Structure comes with maturation. Human Brain Mapping, 32(3), 413–425. 10.1002/hbm.21030

9. Buzsáki, G. (2006). Rhythms of the Brain. Oxford University Press. 10.1093/acprof:oso/9780195301069.001.0001

10. Buzsáki, G., Logothetis, N., & Singer, W. (2013). Scaling Brain Size, Keeping Timing: Evolutionary Preservation of Brain Rhythms. Neuron, 80(3), 751–764. 10.1016/j.neuron.2013.10.002

11. Buzsáki, G., & Watson, B. O. (2012). Brain rhythms and neural syntax: Implications for efficient coding of cognitive content and neuropsychiatric disease. Dialogues in Clinical Neuroscience, 14(4), 345–367. 10.31887/DCNS.2012.14.4/gbuzsaki

12. Capilla, A., Arana, L., García-Huéscar, M., Melcón, M., Gross, J., & Campo, P. (2022). The natural frequencies of the resting human brain: An MEG-based atlas. NeuroImage, 258, 119373. 10.1016/j.neuroimage.2022.119373

13. Chen, A. C. N., Feng, W., Zhao, H., Yin, Y., & Wang, P. (2008). EEG default mode network in the human brain: Spectral regional field powers. NeuroImage, 41(2), 561–574. 10.1016/j.neuroimage.2007.12.064

14. Cho, S., Van Es, M., Woolrich, M., & Gohil, C. (2024). Comparison between EEG and MEG of static and dynamic resting-state networks. Human Brain Mapping, 45(13), e70018. 10.1002/hbm.70018

15. Congedo, M., John, R. E., De Ridder, D., & Prichep, L. (2010). Group independent component analysis of resting state EEG in large normative samples. International Journal of Psychophysiology, 78(2), 89–99. 10.1016/j.ijpsycho.2010.06.003

16. Di Dona, G., & Ronconi, L. (2023). Beta oscillations in vision: A (preconscious) neural mechanism for the dorsal visual stream? Frontiers in Psychology, 14, 1296483. 10.3389/fpsyg.2023.1296483

17. Donoghue, T., Schaworonkow, N., & Voytek, B. (2022). Methodological considerations for studying neural oscillations. European Journal of Neuroscience, 55(11–12), 3502–3527. 10.1111/ejn.15361

18. Ferrarelli, F., Sarasso, S., Guller, Y., Riedner, B. A., Peterson, M. J., Bellesi, M., Massimini, M., Postle, B. R., & Tononi, G. (2012). Reduced Natural Oscillatory Frequency of Frontal Thalamocortical Circuits in Schizophrenia. Archives of General Psychiatry, 69(8). 10.1001/archgenpsychiatry.2012.147

19. Frauscher, B., Von Ellenrieder, N., Zelmann, R., Doležalová, I., Minotti, L., Olivier, A., Hall, J., Hoffmann, D., Nguyen, D. K., Kahane, P., Dubeau, F., & Gotman, J. (2018). Atlas of the normal intracranial electroencephalogram: Neurophysiological awake activity in different cortical areas. Brain, 141(4), 1130–1144. 10.1093/brain/awy035

20. Fries, P. (2015). Rhythms for Cognition: Communication through Coherence. Neuron, 88(1), 220–235. 10.1016/j.neuron.2015.09.034

21. Geller, A. S., Burke, J. F., Sperling, M. R., Sharan, A. D., Litt, B., Baltuch, G. H., Lucas, T. H., & Kahana, M. J. (2014). Eye closure causes widespread low-frequency power increase and focal gamma attenuation in the human electrocorticogram. Clinical Neurophysiology, 125(9), 1764–1773. 10.1016/j.clinph.2014.01.021

22. Getzmann, S., Gajewski, P. D., Schneider, D., & Wascher, E. (2024). Resting-state EEG data before and after cognitive activity across the adult lifespan and a 5-year follow-up. Scientific Data, 11(1), 988. 10.1038/s41597-024-03797-w

23. Gil Ávila, C., Bott, F. S., Tiemann, L., Hohn, V. D., May, E. S., Nickel, M. M., Zebhauser, P. T., Gross, J., & Ploner, M. (2023). DISCOVER-EEG: An open, fully automated EEG pipeline for biomarker discovery in clinical neuroscience. Scientific Data, 10(1), 613. 10.1038/s41597-023-02525-0

24. Goldenholz, D. M., Ahlfors, S. P., Hämäläinen, M. S., Sharon, D., Ishitobi, M., Vaina, L. M., & Stufflebeam, S. M. (2009). Mapping the signal-to-noise-ratios of cortical sources in magnetoencephalography and electroencephalography. Human Brain Mapping, 30(4), 1077–1086. 10.1002/hbm.20571

25. Groppe, D. M., Bickel, S., Keller, C. J., Jain, S. K., Hwang, S. T., Harden, C., & Mehta, A. D. (2013). Dominant frequencies of resting human brain activity as measured by the electrocorticogram. NeuroImage, 79, 223–233. 10.1016/j.neuroimage.2013.04.044

26. Haegens, S., Cousijn, H., Wallis, G., Harrison, P. J., & Nobre, A. C. (2014). Inter- and intra-individual variability in alpha peak frequency. NeuroImage, 92, 46–55. 10.1016/j.neuroimage.2014.01.049

27. Herrera-Morueco, J. J., Stern, E., Arana, L., Capilla, A. (2026). Globally stable, locally flexible: Dynamic reconfiguration of brain natural frequencies during cognitive processing. BioRxiv

28. Hillebrand, A., Barnes, G. R., Bosboom, J. L., Berendse, H. W., & Stam, C. J. (2012). Frequency-dependent functional connectivity within resting-state networks: An atlas-based MEG beamformer solution. NeuroImage, 59(4), 3909–3921. 10.1016/j.neuroimage.2011.11.005

29. Holmes, C. J., Hoge, R., Collins, L., Woods, R., Toga, A. W., & Evans, A. C. (1998). Enhancement of MR images using registration for signal averaging. Journal of Computer Assisted Tomography, 22(2), 324–333. 10.1097/00004728-199803000-00032

30. Janiukstyte, V., Owen, T. W., Chaudhary, U. J., Diehl, B., Lemieux, L., Duncan, J. S., De Tisi, J., Wang, Y., & Taylor, P. N. (2023). Normative brain mapping using scalp EEG and potential clinical application. Scientific Reports, 13(1), 13442. 10.1038/s41598-023-39700-7

31. Kalamangalam, G. P., Long, S., & Chelaru, M. I. (2020). A neurophysiological brain map: Spectral parameterization of the human intracranial electroencephalogram. Clinical Neurophysiology, 131(3), 665–675. 10.1016/j.clinph.2019.11.061

32. Keitel, A., & Gross, J. (2016). Individual Human Brain Areas Can Be Identified from Their Characteristic Spectral Activation Fingerprints. PLOS Biology, 14(6), e1002498. 10.1371/journal.pbio.1002498

33. Klamer, S., Elshahabi, A., Lerche, H., Braun, C., Erb, M., Scheffler, K., & Focke, N. K. (2015). Differences Between MEG and High-Density EEG Source Localizations Using a Distributed Source Model in Comparison to fMRI. Brain Topography, 28(1), 87–94. 10.1007/s10548-014-0405-3

34. Klem, G. H., Lüders, H. O., Jasper, H., & Elger, C. (1999). The ten-twenty electrode system of the International Federation. The International Federation of Clinical Neurophysiology. *Electroencephalography and Clinical Neurophysiology*, Supplement, 52, 3–6.

35. Leske, S., & Dalal, S. S. (2019). Reducing power line noise in EEG and MEG data via spectrum interpolation. NeuroImage, 189, 763–776. 10.1016/j.neuroimage.2019.01.026

36. Lew, B. J., Fitzgerald, E. E., Ott, L. R., Penhale, S. H., & Wilson, T. W. (2021). Three-year reliability of MEG resting-state oscillatory power. NeuroImage, 243, 118516. 10.1016/j.neuroimage.2021.118516

37. Liu, A. K., Dale, A. M., & Belliveau, J. W. (2002). Monte Carlo simulation studies of EEG and MEG localization accuracy. Human Brain Mapping, 16(1), 47–62. 10.1002/hbm.10024

38. Lopes da Silva, F. (2013). EEG and MEG: Relevance to Neuroscience. Neuron, 80(5), 1112–1128. 10.1016/j.neuron.2013.10.017

39. Mahjoory, K., Schoffelen, J.-M., Keitel, A., & Gross, J. (2020). The frequency gradient of human resting-state brain oscillations follows cortical hierarchies. eLife, 9, e53715. 10.7554/eLife.53715

40. Malmivuo, J. (2012). Comparison of the Properties of EEG and MEG in Detecting the Electric Activity of the Brain. Brain Topography, 25(1), 1–19. 10.1007/s10548-011-0202-1

41. Mellem, M. S., Wohltjen, S., Gotts, S. J., Ghuman, A. S., & Martin, A. (2017). Intrinsic frequency biases and profiles across human cortex. Journal of Neurophysiology, 118(5), 2853–2864. 10.1152/jn.00061.2017

42. Mierau, A., Klimesch, W., & Lefebvre, J. (2017). State-dependent alpha peak frequency shifts: Experimental evidence, potential mechanisms and functional implications. Neuroscience, 360, 146–154. 10.1016/j.neuroscience.2017.07.037

43. Mullen, T. R., Kothe, C. A. E., Chi, Y. M., Ojeda, A., Kerth, T., Makeig, S., Jung, T.-P., & Cauwenberghs, G. (2015). Real-time neuroimaging and cognitive monitoring using wearable dry EEG. IEEE Transactions on Biomedical Engineering, 62(11), 2553–2567. 10.1109/TBME.2015.2481482

44. Niso, G., Rogers, C., Moreau, J. T., Chen, L.-Y., Madjar, C., Das, S., Bock, E., Tadel, F., Evans, A. C., Jolicoeur, P., & Baillet, S. (2016). OMEGA: The Open MEG Archive. NeuroImage, 124, 1182–1187. 10.1016/j.neuroimage.2015.04.028

45. Niso, G., Tadel, F., Bock, E., Cousineau, M., Santos, A., & Baillet, S. (2019). Brainstorm Pipeline Analysis of Resting-State Data From the Open MEG Archive. Frontiers in Neuroscience, 13, 284. 10.3389/fnins.2019.00284

46. Nolte, G. (2003). The magnetic lead field theorem in the quasi-static approximation and its use for magnetoencephalography forward calculation in realistic volume conductors. Physics in Medicine and Biology, 48(22), 3637–3652. 10.1088/0031-9155/48/22/002

47. Oostenveld, R., Fries, P., Maris, E., & Schoffelen, J.-M. (2011). FieldTrip: Open Source Software for Advanced Analysis of MEG, EEG, and Invasive Electrophysiological Data. Computational Intelligence and Neuroscience, 2011, 1–9. 10.1155/2011/156869

48. Oostenveld, R., & Praamstra, P. (2001). The fivepercent electrode system for high-resolution EEG and ERP measurements. 112, 713–719.

49. Oostenveld, R., Stegeman, D. F., Praamstra, P., & Van Oosterom, A. (2003). Brain symmetry and topographic analysis of lateralized event-related potentials. Clinical Neurophysiology, 114(7), 1194–1202. 10.1016/S1388-2457(03)00059-2

50. Pernet, C. R., Martinez-Cancino, R., Truong, D., Makeig, S., & Delorme, A. (2021). From BIDS-Formatted EEG Data to Sensor-Space Group Results: A Fully Reproducible Workflow With EEGLAB and LIMO EEG. Frontiers in Neuroscience, 14, 610388. 10.3389/fnins.2020.610388

51. Perrin, F., Pernier, J., Bertrand, O., & Echallier, J. F. (1989). Spherical splines for scalp potential and current density mapping. Electroencephalography and Clinical Neurophysiology, 72(2), 184–187. 10.1016/0013-4694(89)90180-6

52. Petro, N. M., Ott, L. R., Penhale, S. H., Rempe, M. P., Embury, C. M., Picci, G., Wang, Y.-P., Stephen, J. M., Calhoun, V. D., & Wilson, T. W. (2022). Eyes-closed versus eyes-open differences in spontaneous neural dynamics during development. NeuroImage, 258, 119337. 10.1016/j.neuroimage.2022.119337

53. Pfurtscheller, G., & Cooper, R. (1975). Frequency dependence of the transmission of the EEG from cortex to scalp. Electroencephalography and Clinical Neurophysiology, 38(1), 93–96. 10.1016/0013-4694(75)90215-1

54. Pion-Tonachini, L., Kreutz-Delgado, K., & Makeig, S. (2019). ICLabel: An automated electroencephalographic independent component classifier, dataset, and website. NeuroImage, 198, 181–197. 10.1016/j.neuroimage.2019.05.026

55. Quigley, C. (2022). Forgotten rhythms? Revisiting the first evidence for rhythms in cognition. European Journal of Neuroscience, 55(11–12), 3266–3276. 10.1111/ejn.15450

56. Ramkumar, P., Parkkonen, L., & Hyvärinen, A. (2014). Group-level spatial independent component analysis of Fourier envelopes of resting-state MEG data. NeuroImage, 86, 480–491. 10.1016/j.neuroimage.2013.10.032

57. Rosanova, M., Casali, A., Bellina, V., Resta, F., Mariotti, M., & Massimini, M. (2009). Natural Frequencies of Human Corticothalamic Circuits. Journal of Neuroscience, 29(24), 7679–7685. 10.1523/JNEUROSCI.0445-09.2009

58. Salmelin, R., & Hari, R. (1994). Characterization of spontaneous MEG rhythms in healthy adults. Electroencephalography and Clinical Neurophysiology, 91(4), 237–248. 10.1016/0013-4694(94)90187-2

59. Schnitzler, A., & Gross, J. (2005). Normal and pathological oscillatory communication in the brain. Nature Reviews Neuroscience, 6(4), 285–296. 10.1038/nrn1650

60. Shapira Lots, I., Zeev-Wolf, M., Harpaz, Y., & Abeles, M. (2016). Source localization scale correction for Beamformer analysis. Journal of Neuroscience Methods, 273, 10–19. 10.1016/j.jneumeth.2016.07.013

61. Srinivasan, R., Winter, W. R., & Nunez, P. L. (2006). Source analysis of EEG oscillations using high-resolution EEG and MEG. In Progress in Brain Research (Vol. 159, pp. 29–42). Elsevier. 10.1016/S0079-6123(06)59003-X

62. Takeuchi, Y., & Berényi, A. (2020). Oscillotherapeutics – Time-targeted interventions in epilepsy and beyond. Neuroscience Research, 152, 87–107. 10.1016/j.neures.2020.01.002

63. Van Veen, B. D., Van Drongelen, W., Yuchtman, M., & Suzuki, A. (1997). Localization of brain electrical activity via linearly constrained minimum variance spatial filtering. IEEE Transactions on Biomedical Engineering, 44(9), 867–880. 10.1109/10.623056

64. Varela, F., Lachaux, J.-P., Rodriguez, E., & Martinerie, J. (2001). The brainweb: Phase synchronization and large-scale integration. Nature Reviews Neuroscience, 2(4), 229–239. 10.1038/35067550

